## Supplementary Materials for "MaCroDNA: Accurate integration of single-cell DNA and RNA data for a deeper understanding of tumor heterogeneity"

### 1 Supplementary figures

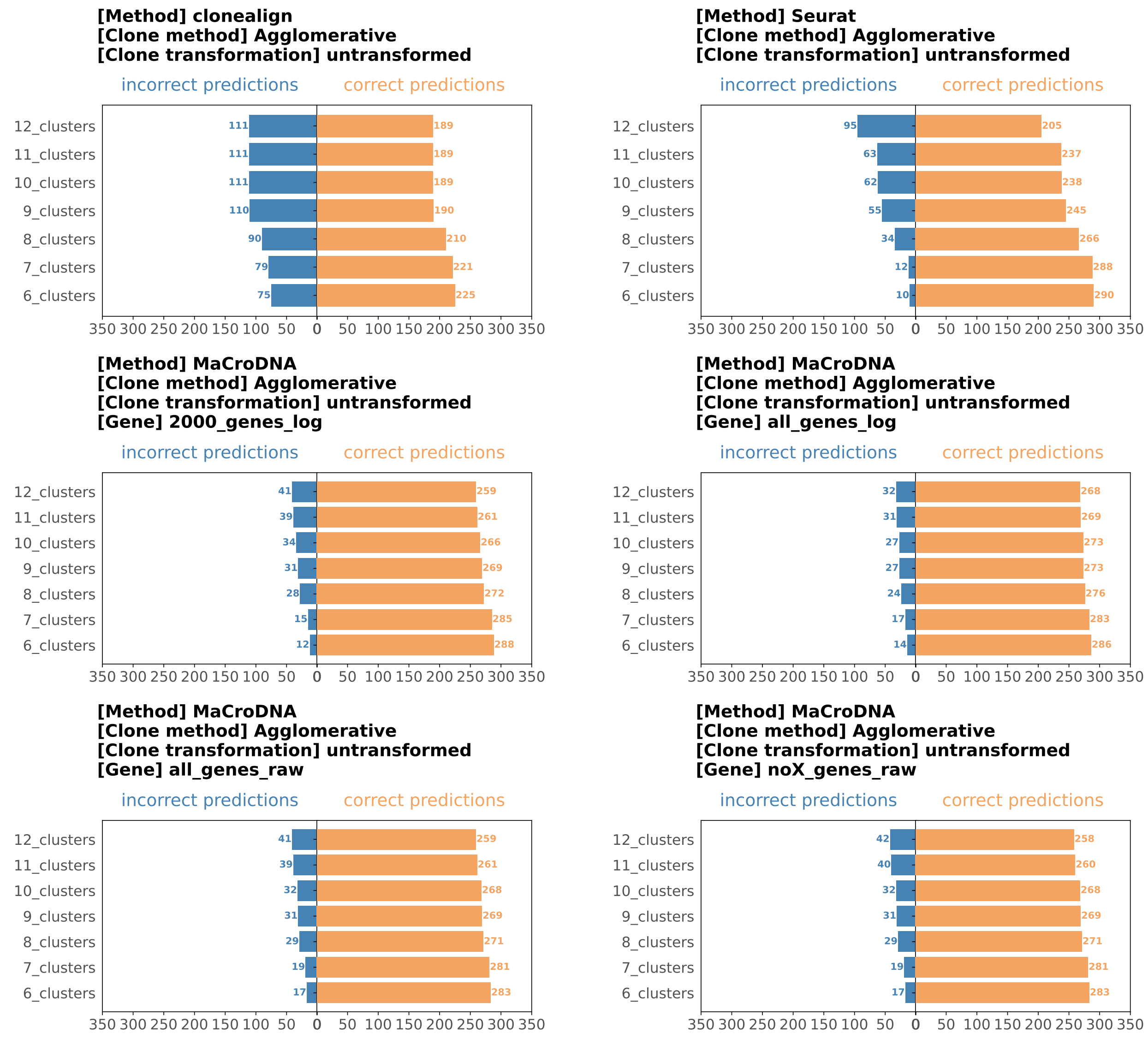


Figure S1: **Results of agglomerative clustering method, using untransformed data for clustering.** The input of clonealign is the original data without genes on X-chromosome. The input of Seurat is the log-transformed data with the top 2000 genes having been selected. The input of MaCroDNA has four different settings: `2000_genes_log` is same as the input of Seurat, `all_genes_log` uses the same log-transformation as Seurat but all genes are kept, `all_genes_raw` is the original data with all genes, and `noX_genes_raw` is same as the input of clonealign, where the genes on X-chromosome are removed and the values are the original values.

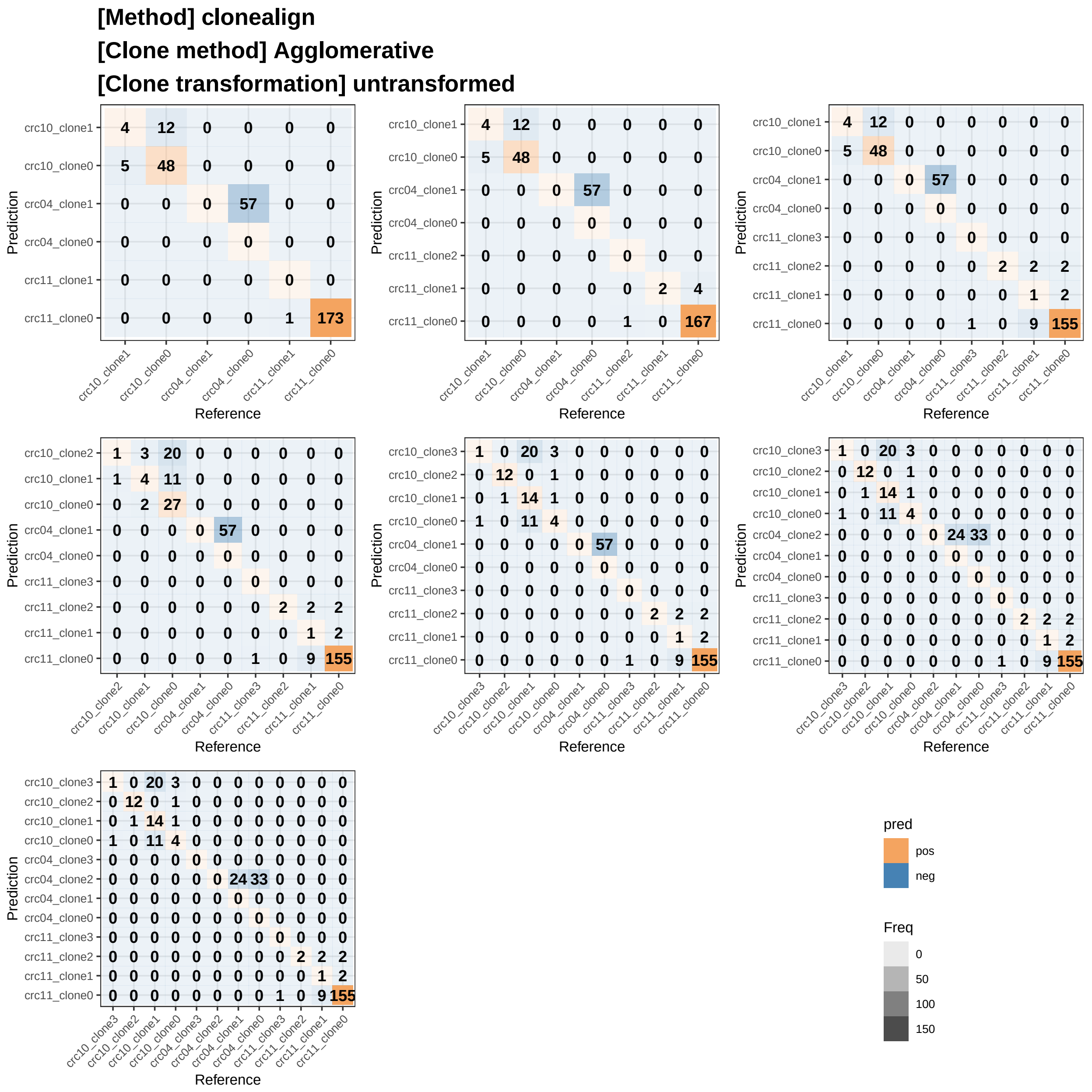


Figure S2: **clonealign results using agglomerative clustering method and the original data for clustering.** The input for the model is the original value without the genes on X-chromosome. The data for clustering is the original untransformed data.

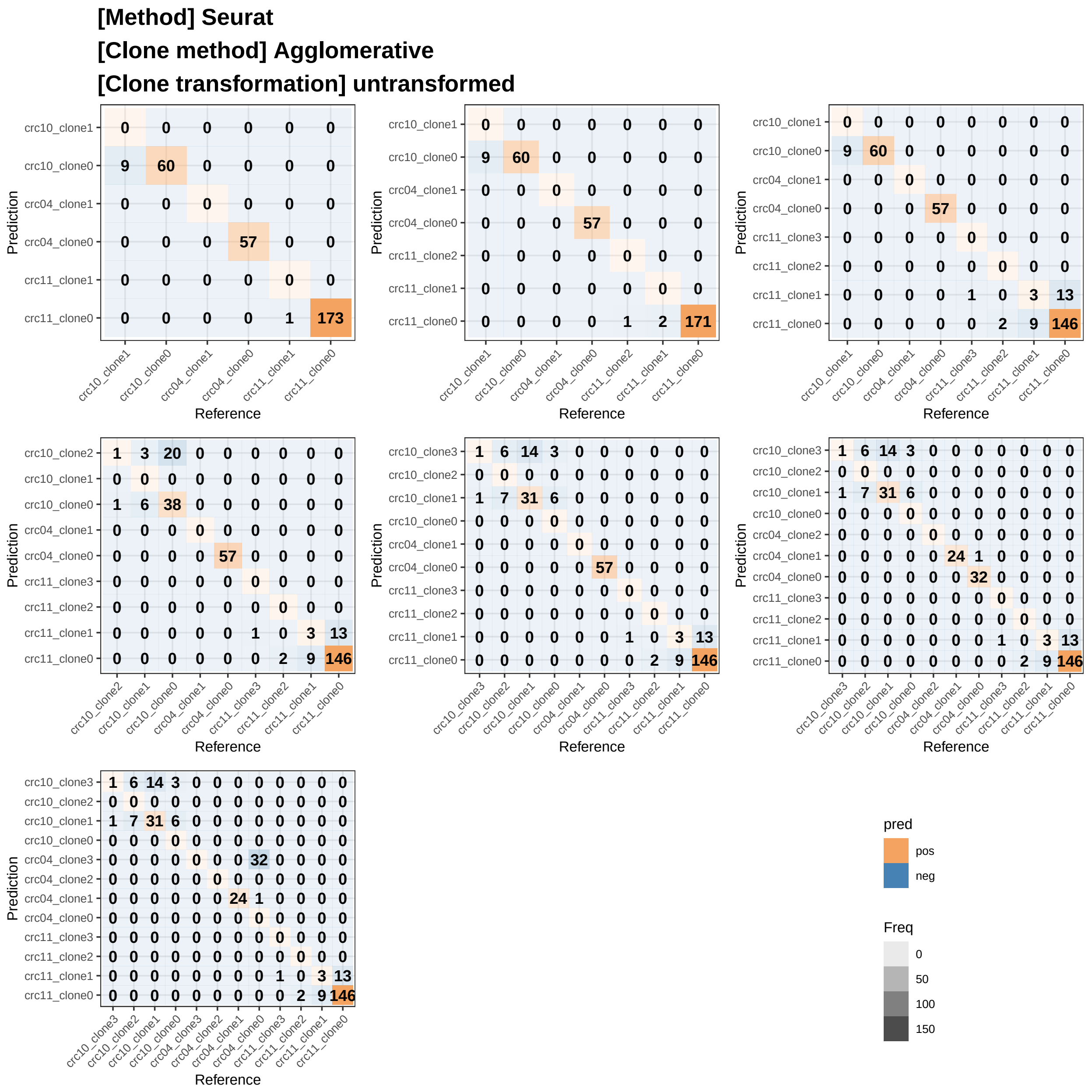


Figure S3: **Seurat results using agglomerative clustering method and the original data for clustering.** The input for the model is the log-transformed data of the original data with the top 2000 genes having been selected. The data for clustering is the original data.

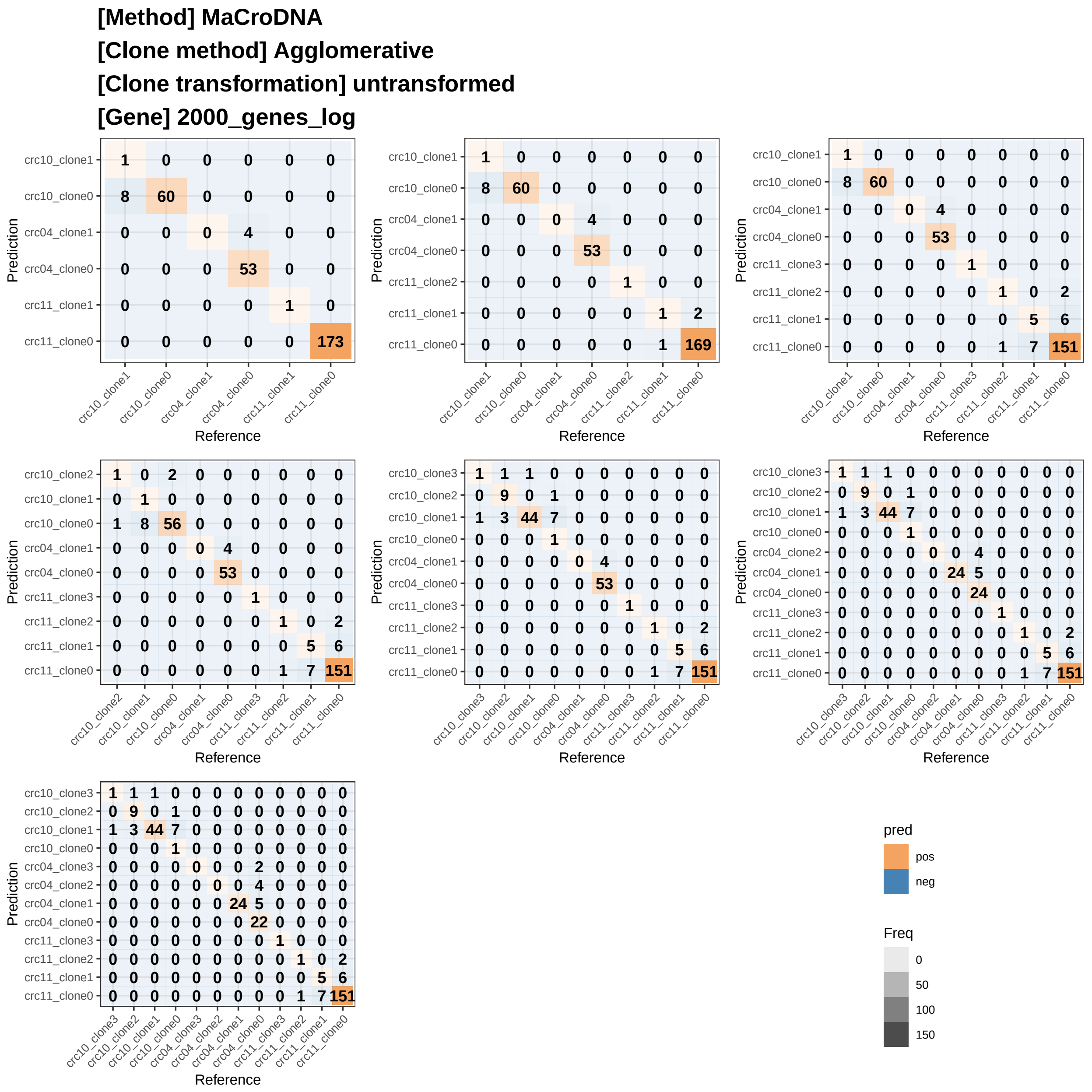


Figure S4: **MaCroDNA results using agglomerative clustering method and the original data for clustering.** The input for the model is same as the input for Seurat. The input is the log-transformed data of the original data with the top 2000 genes having been selected. The data for clustering is the original data.

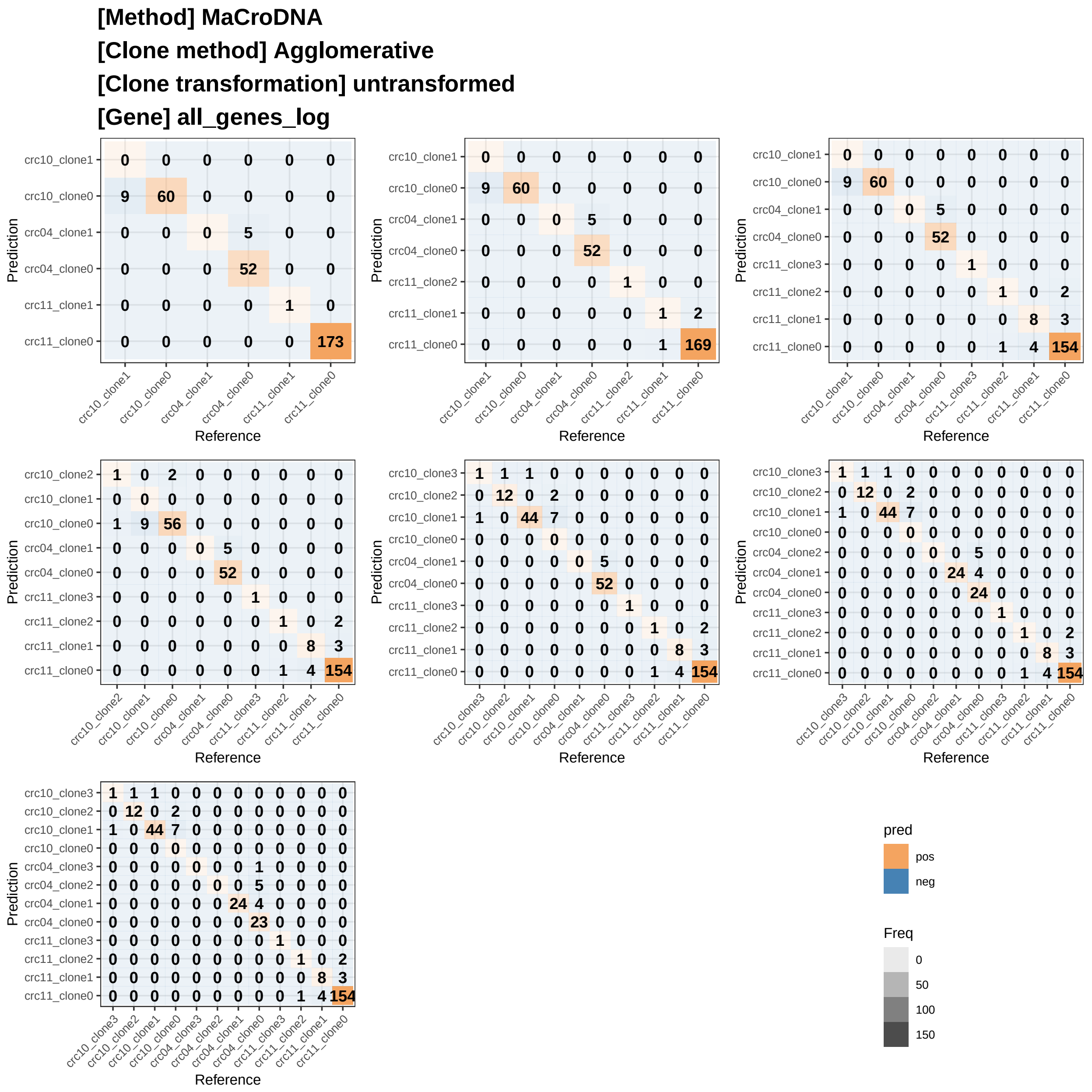


Figure S5: **MaCroDNA results using agglomerative clustering method and the original data for clustering.** The input for the model is the log-transformed data of the original data, same transformation as the input for Seurat. But all the genes are used for the model input. The data for clustering is the original data.

**[Method] MaCroDNA**
**[Clone method] Agglomerative**
**[Clone transformation] untransformed**
**[Gene] all_genes_raw**

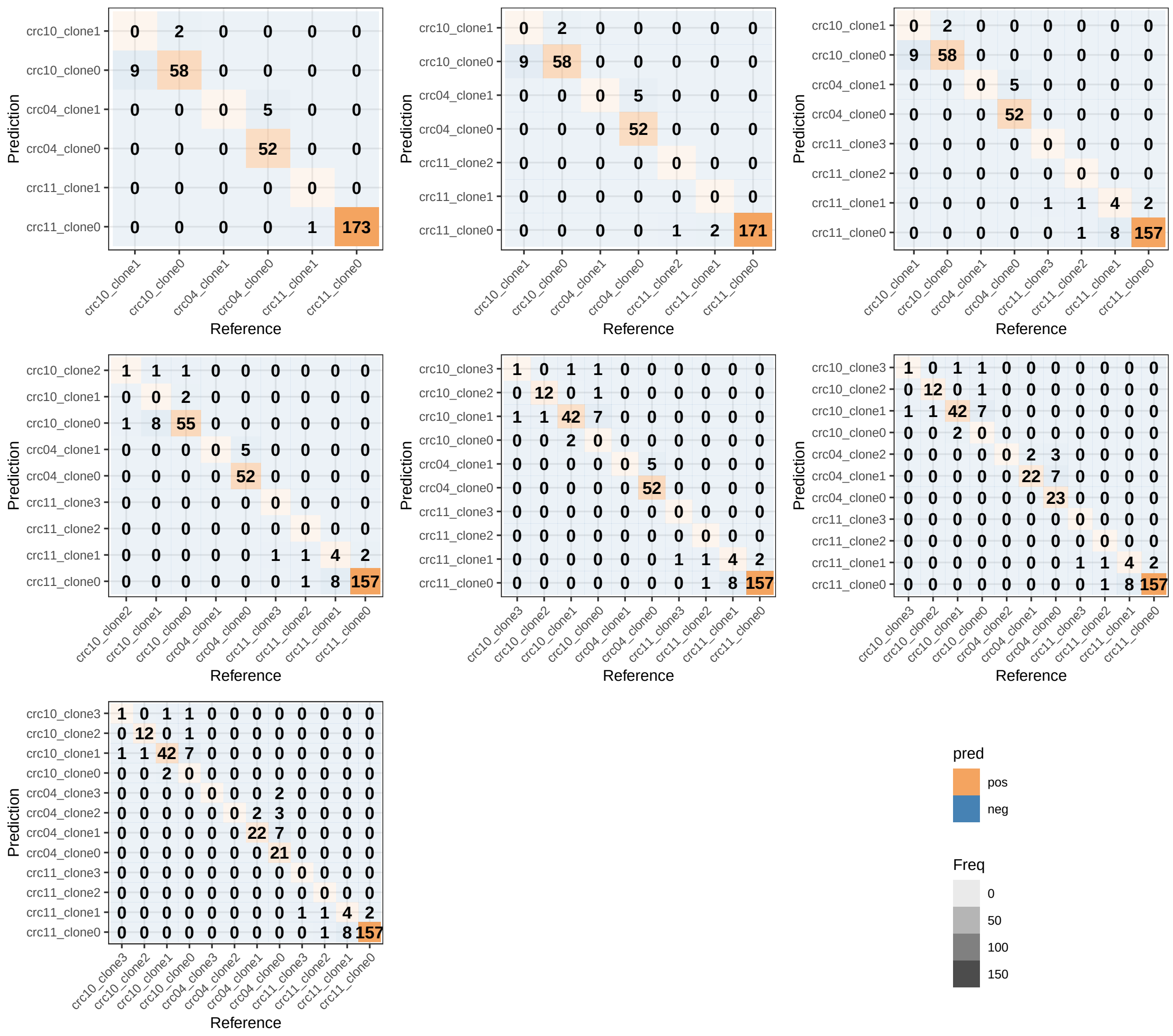

Figure S6: **MaCroDNA results using agglomerative clustering method and the original data for clustering.** The input for the model is the original data with all the genes. The data for clustering is the original data.

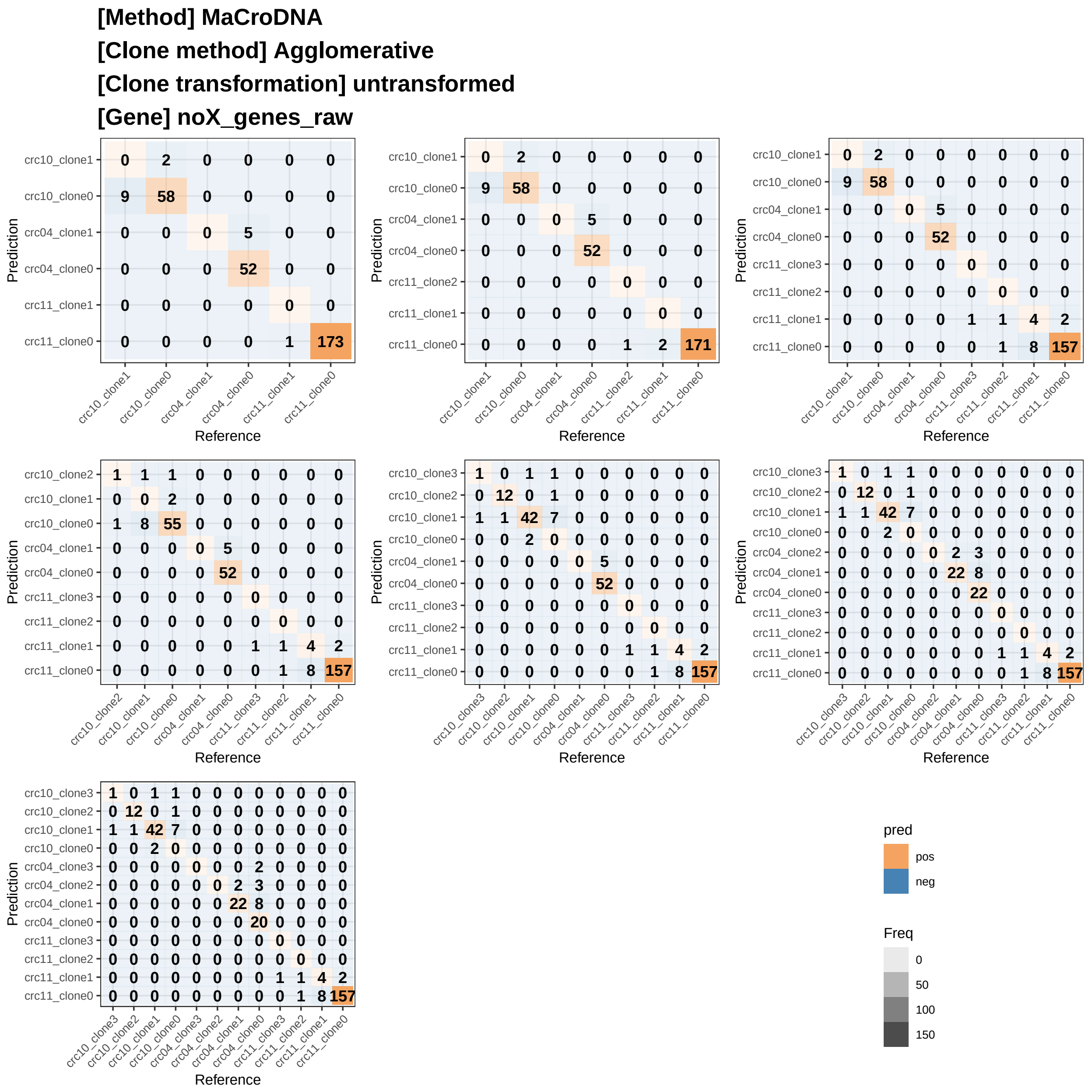


Figure S7: **MaCroDNA results using agglomerative clustering method and the original data for clustering.** The input for the model is the original data without the genes on X-chromosome, same as the input for clonealign. The data for clustering is the original data.

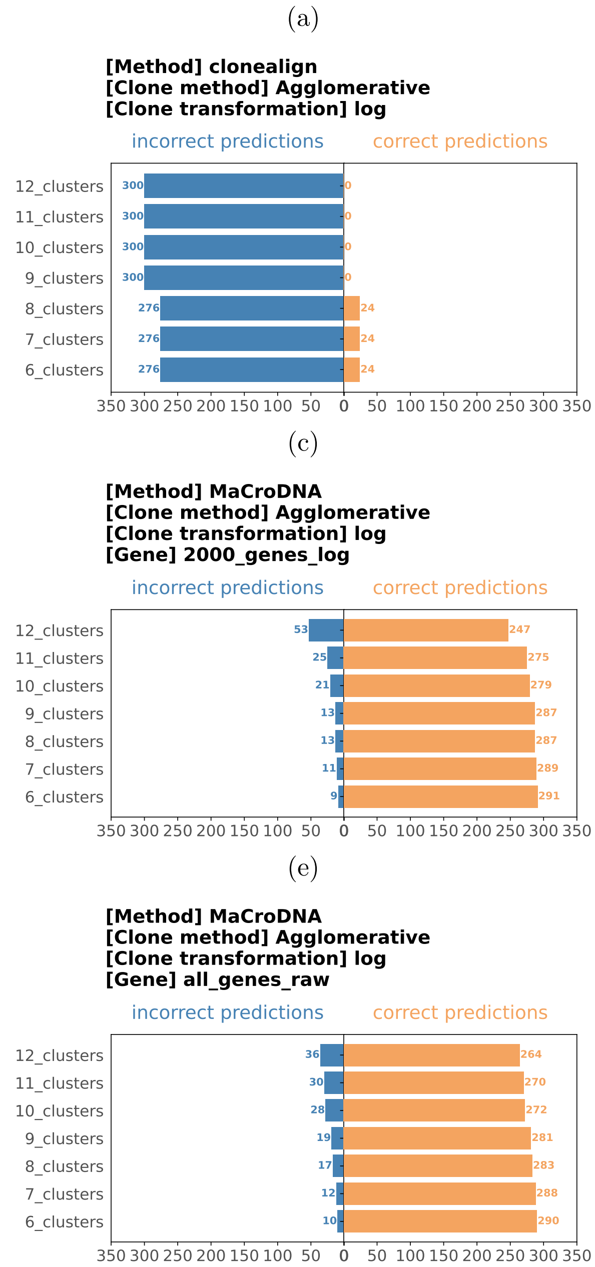


Figure S8: **Results of agglomerative clustering method, using log-transformed data for clustering.** The input of clonealign is the original data without genes on X-chromosome. The input of Seurat is the log-transformed data with the top 2000 genes having been selected. The input of MaCroDNA has four different settings: `2000_genes_log` is same as the input of Seurat, `all_genes_log` uses the same log-transformation as Seurat but all genes are used, `all_genes_raw` is the original data with all genes, and `noX_genes_raw` is same as the input of clonealign, where the genes on X-chromosome are removed and the values are the original values.

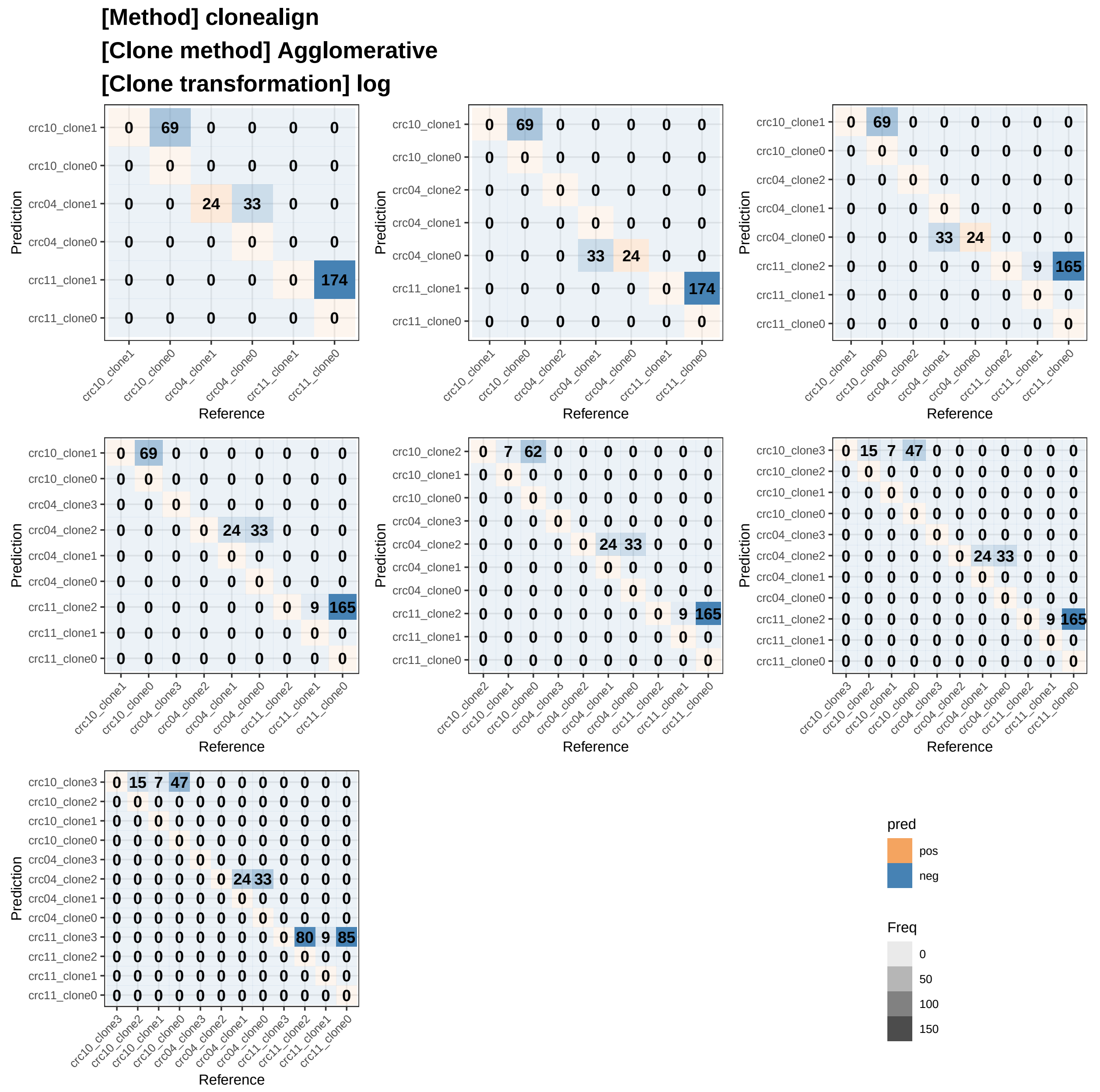


Figure S9: **clonealign results using agglomerative clustering method and the log-transformed data for clustering.**. The input for the model is the original value without the genes on X-chromosome, and the data for clustering is the original log-transformed data.

**[Method] Seurat**

**[Clone method] Agglomerative**

**[Clone transformation] log**

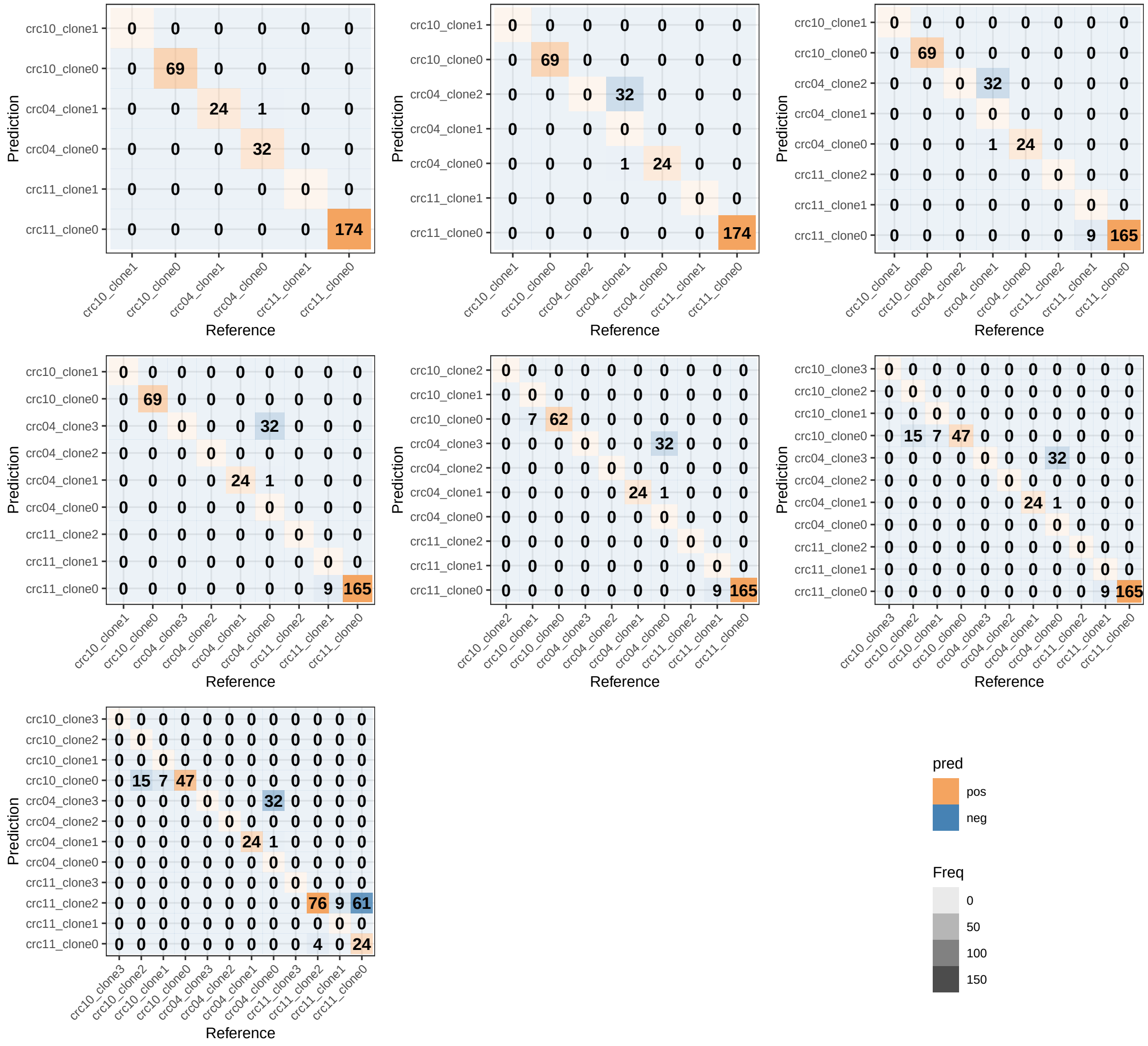


Figure S10: **Seurat results using agglomerative clustering method and the log-transformed data for clustering.** The input for the model is the log-transformed data of the original data with the top 2000 genes having been selected. The data for clustering is the log-transformed data.

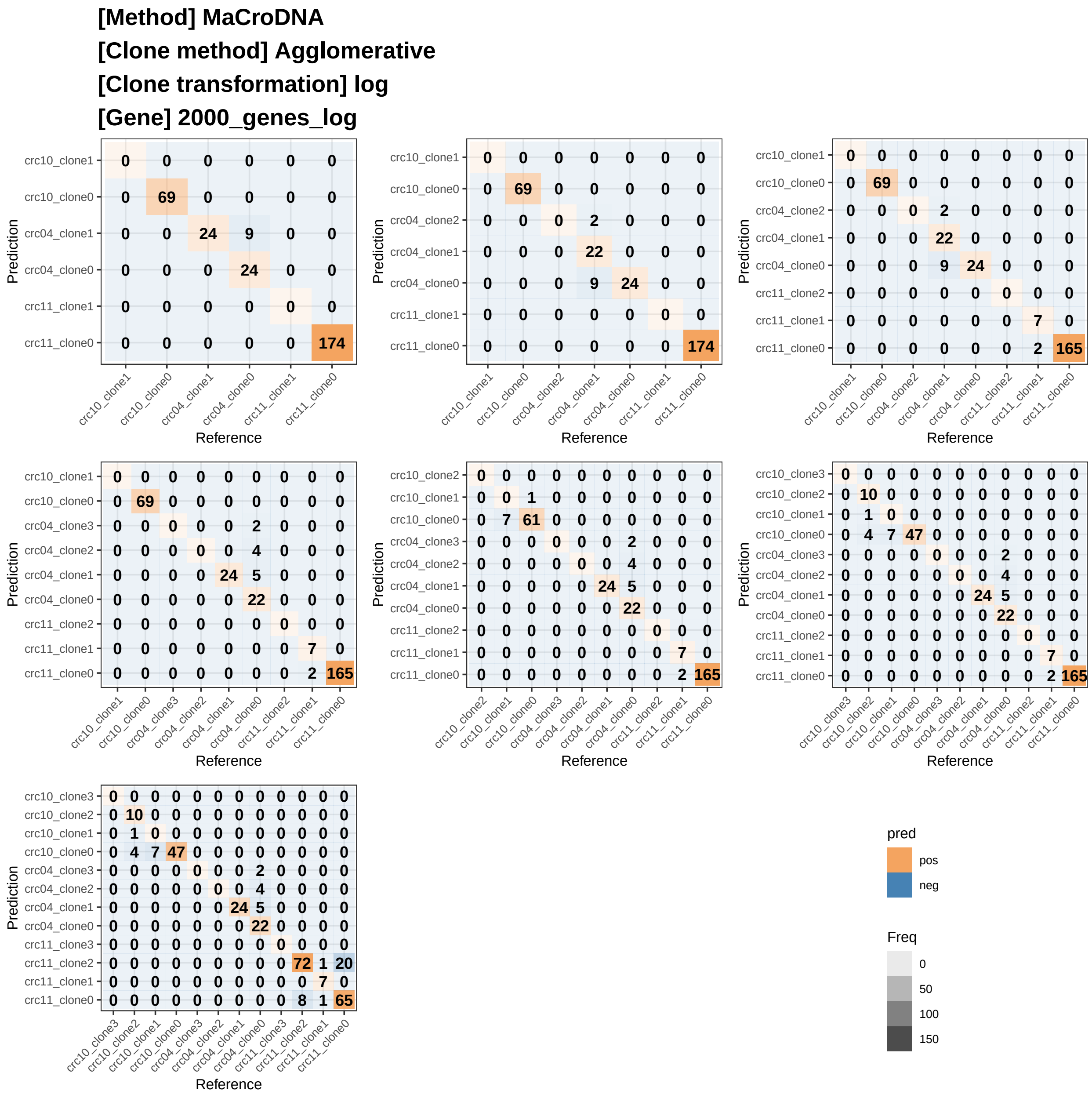


Figure S11: **MaCroDNA results using agglomerative clustering method and the log-transformed data for clustering.** The input for the model is same as the input for Seurat. The input is the log-transformed data of the original data. The top 2000 genes have been selected. The data for clustering is the log-transformed data.

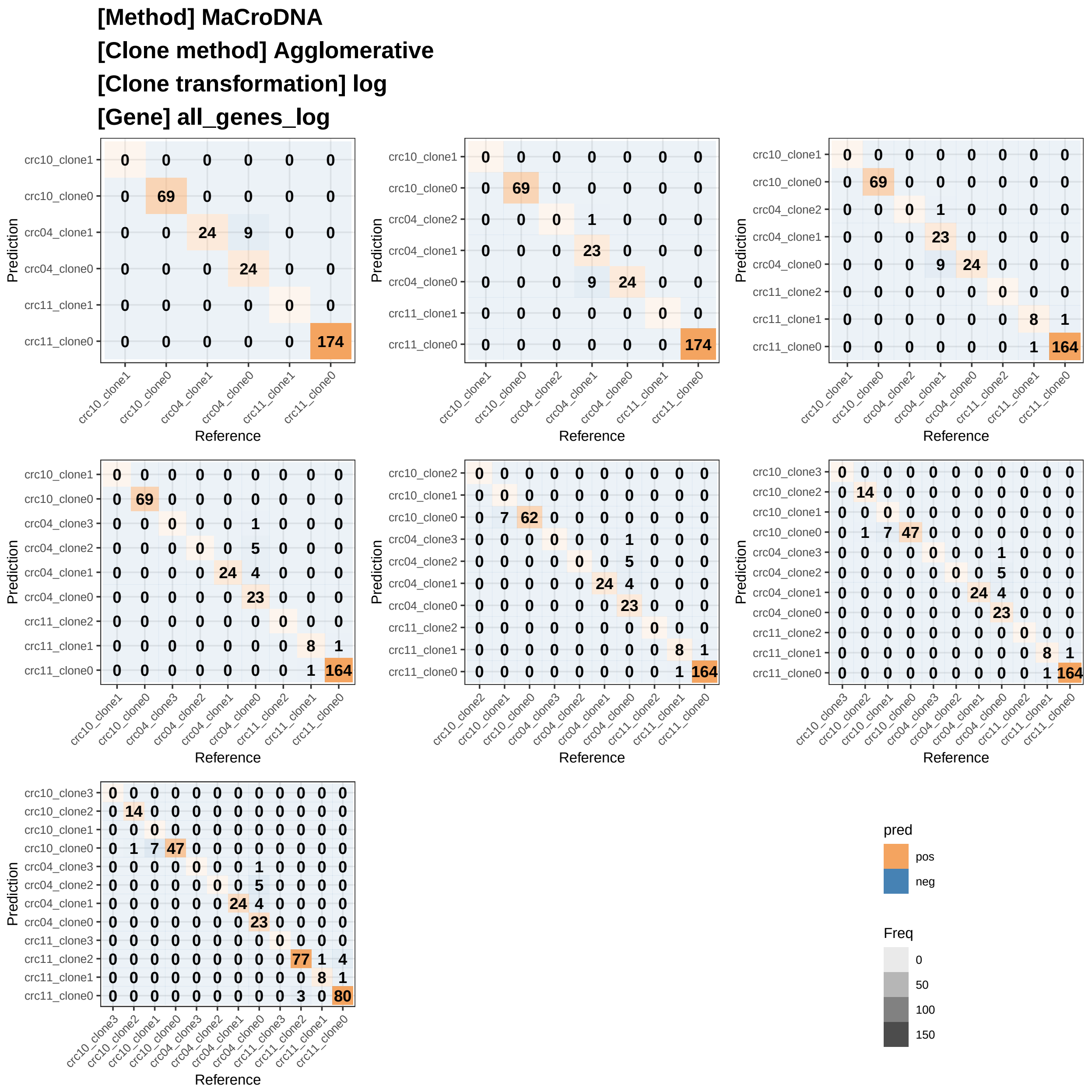


Figure S12: **MaCroDNA results using agglomerative clustering method and the log-transformed data for clustering.** The input for the model is the log-transformed data of the original data, same transformation as the input for Seurat. But all the genes are used as the model input. The data for clustering is the log-transformed data.

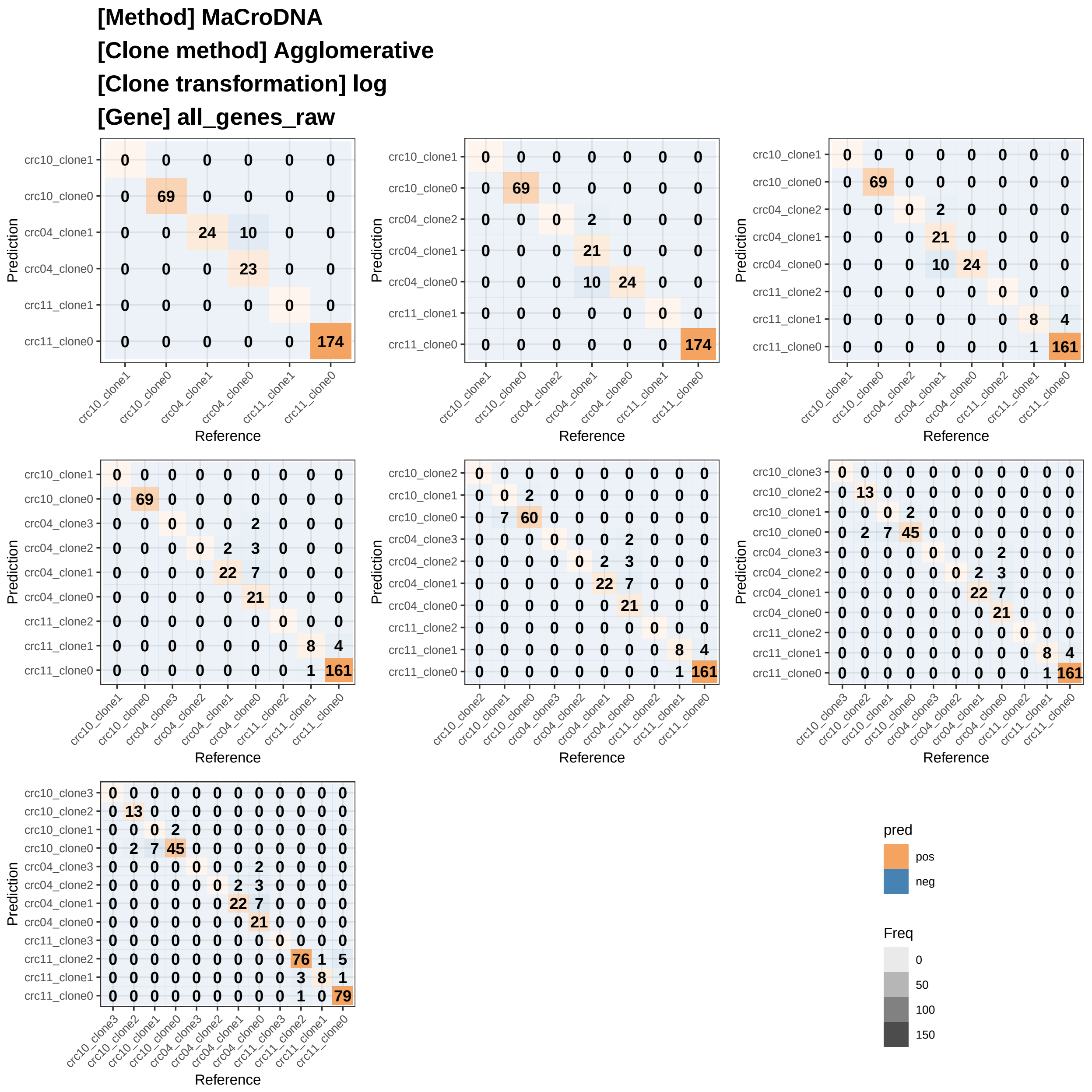


Figure S13: **MaCroDNA results using agglomerative clustering method and the log-transformed data for clustering.** The input for the model is the original data with all the genes. The data for clustering is the log-transformed data.

**[Method] MaCroDNA**
**[Clone method] Agglomerative**
**[Clone transformation] log**
**[Gene] noX_genes_raw**

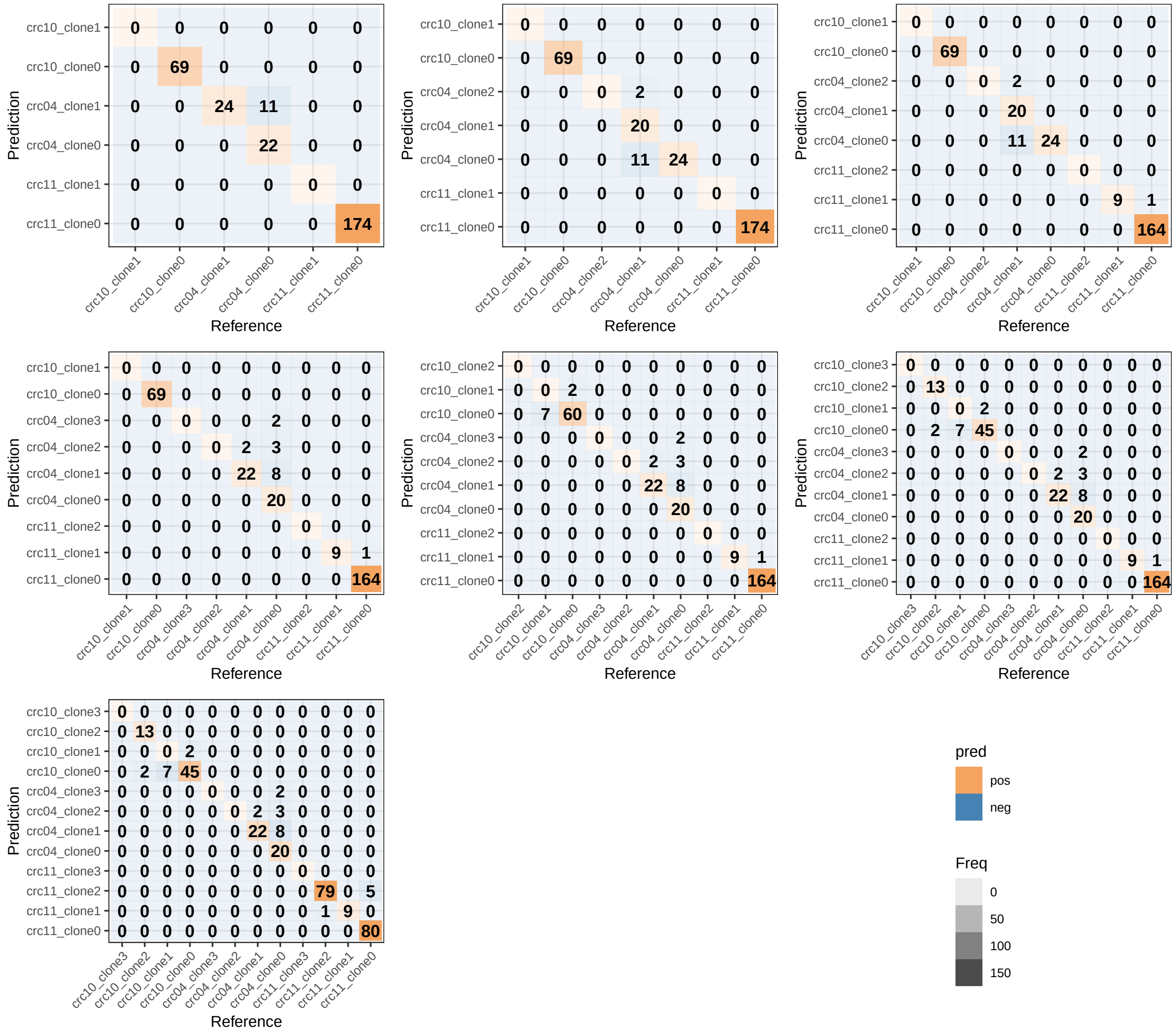


Figure S14: **MaCroDNA results using agglomerative clustering method and the log-transformed data for clustering.** The input for the model is the original data without the genes on X-chromosome, same as the input for clonealign. The data for clustering is the log-transformed data.

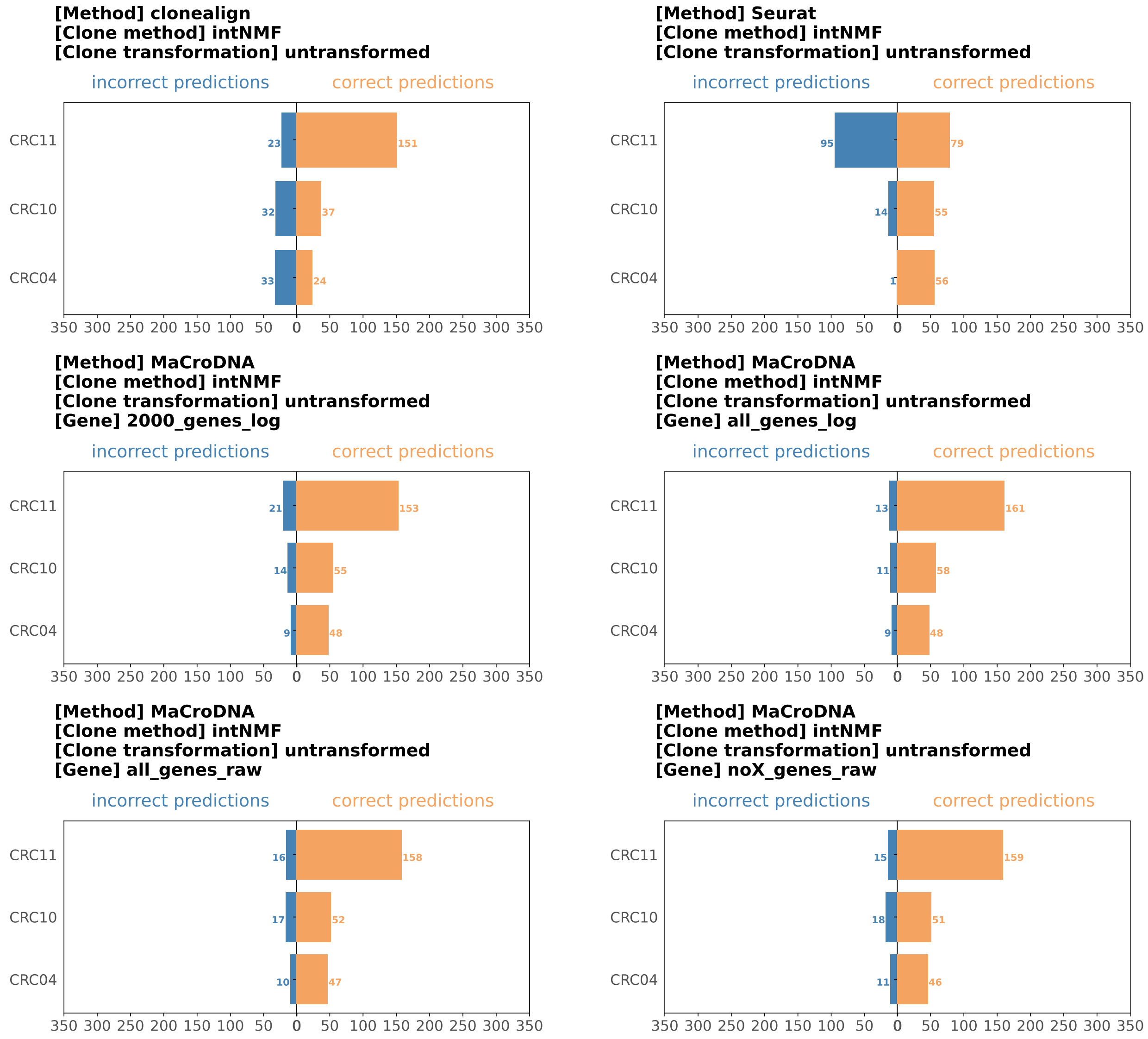


Figure S15: **Results of intNMF clustering method, using untransformed data for clustering.** The input of clonealign is the original data without genes on X-chromosome. The input of Seurat is the log-transformed data with the top 2000 genes having been selected. The input of MaCroDNA has four different settings. `2000_genes_log` is same as the input of Seurat, `all_genes_log` uses the same log-transformation for Seurat but all genes are used, `all_genes_raw` is the original data with all genes, and `noX_genes_raw` is same as the input of clonealign, where the genes on X-chromosome are removed and the values are the original values.

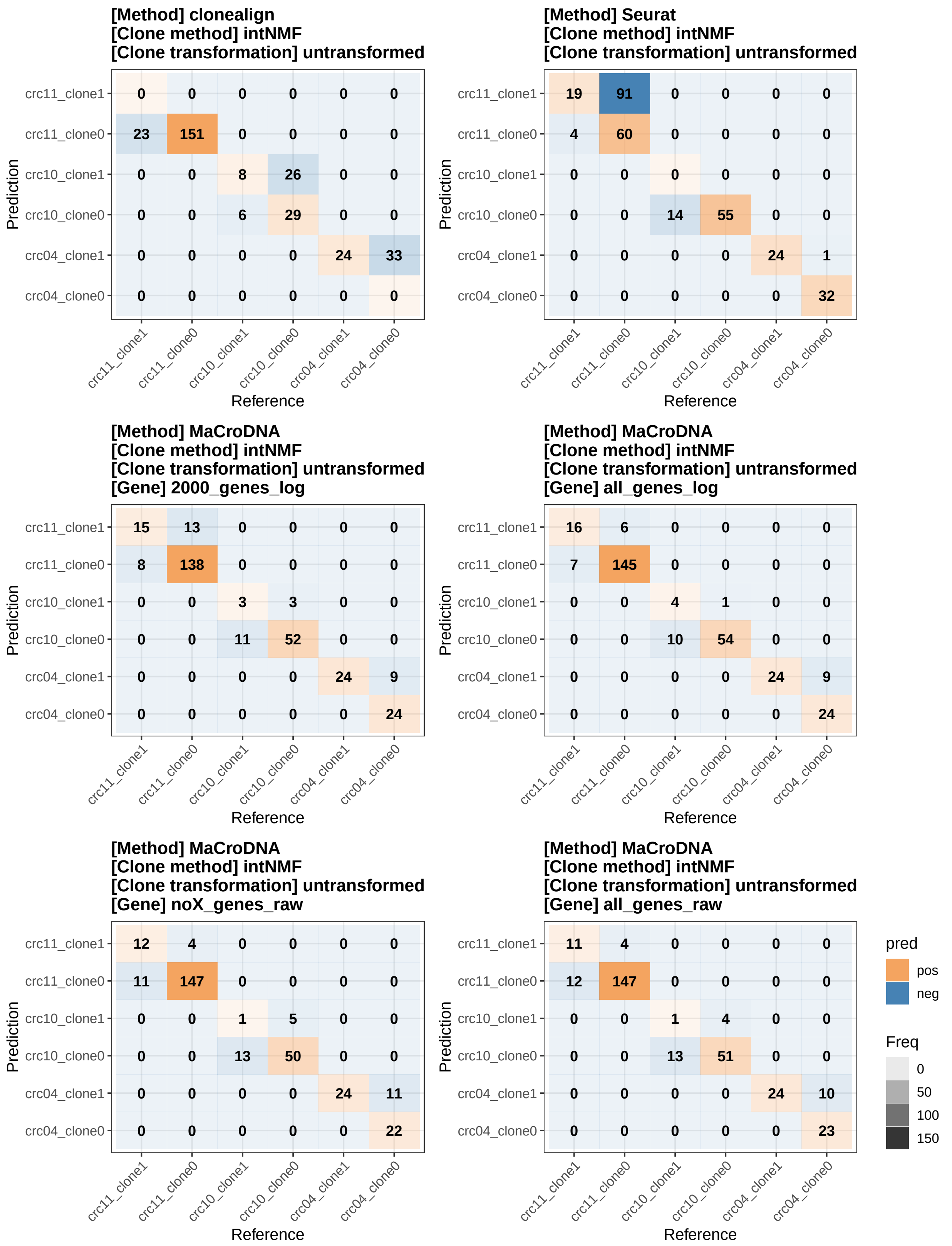


Figure S16: **Results of intNMF clustering method, using untransformed data for clustering.** The input of clonealign is the original data without genes on X-chromosome. The input of Seurat is the log-transformed data with the top 2000 genes having been selected. The input of MaCroDNA has four different settings: `2000_genes_log` is same as the input of Seurat, `all_genes_log` uses the same log-transformation for Seurat but all genes are used, `all_genes_raw` is the original data with all genes, and `noX_genes_raw` is same as the input of clonealign, where the genes on X-chromosome are removed and the values are the original values.

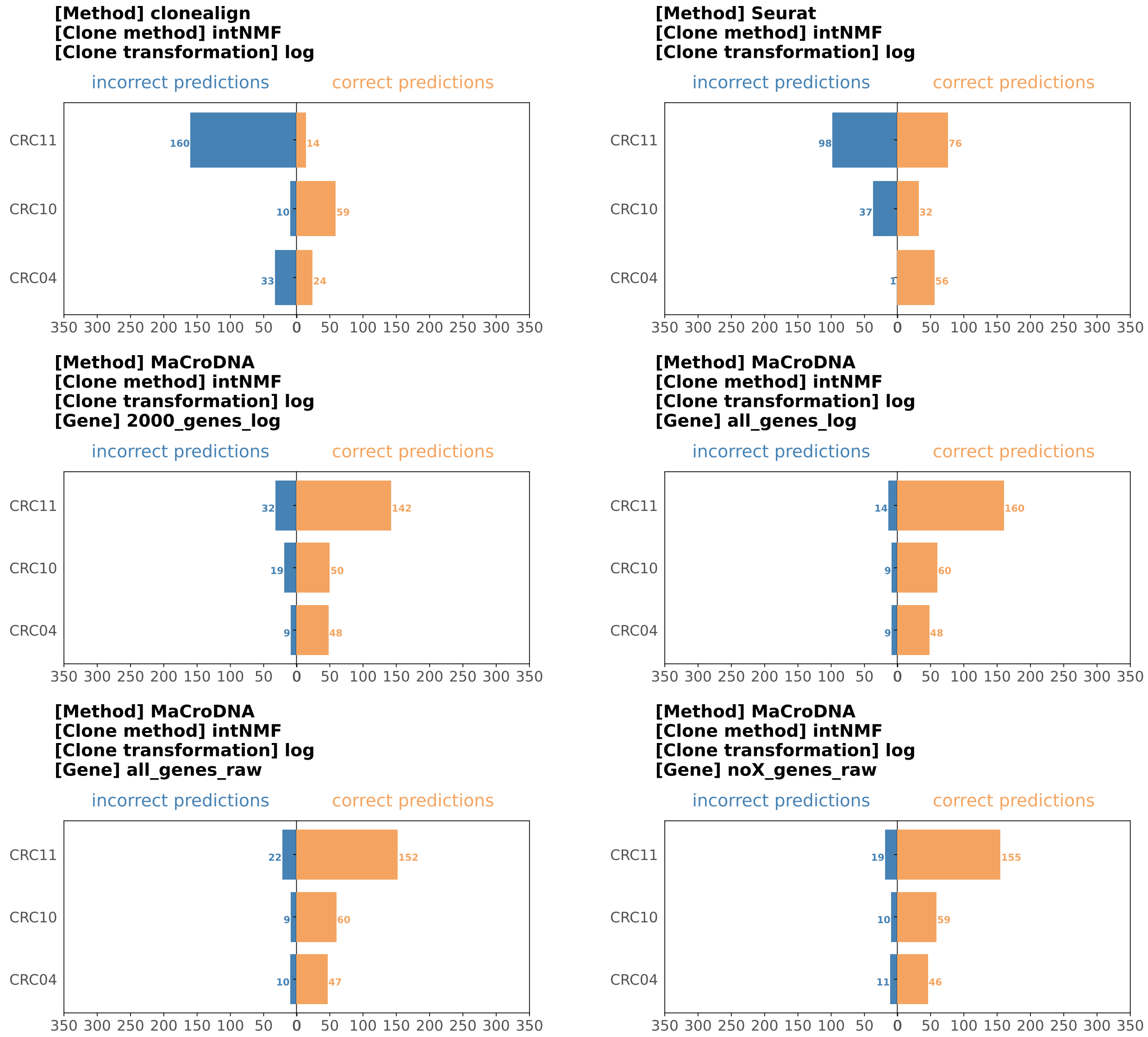


Figure S17: **Results of intNMF clustering method, using log-transformed data for clustering.** The input of clonealign is the original data without genes on X-chromosome. The input of Seurat is the log-transformed data with the top 2000 genes having been selected. The input of MaCroDNA has four different settings: `2000_genes_log` is same as the input of Seurat, `all_genes_log` uses the same log-transformation for Seurat but all genes are used, `all_genes_raw` is the original data with all genes, and `noX_genes_raw` is same as the input of clonealign, where the genes on X-chromosome are removed and the values are the original values.

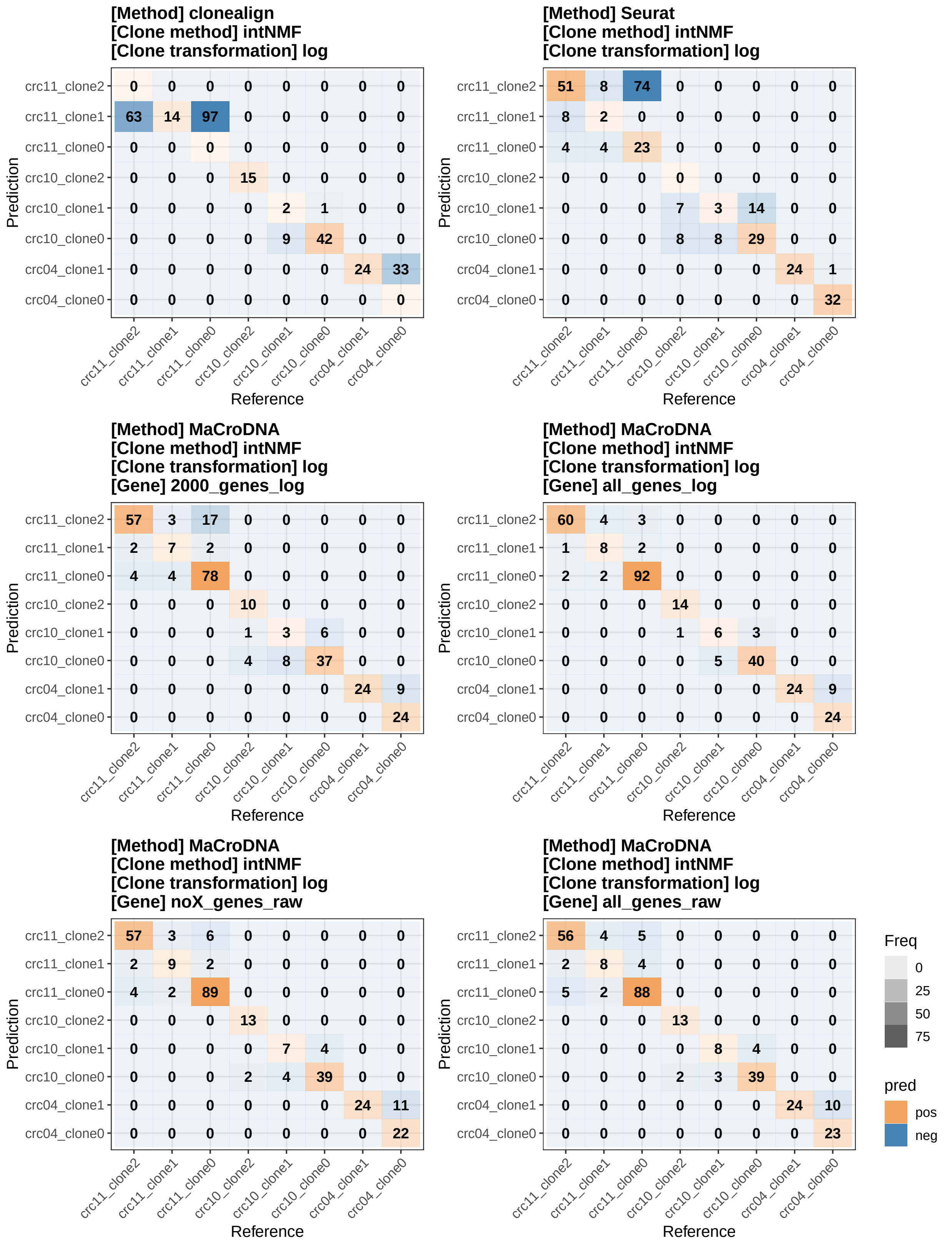


Figure S18: **Results of intNMF clustering method, using log-transformed data for clustering.** The input of clonealign is the original data without genes on X-chromosome. The input of Seurat is the log-transformed data with the top 2000 genes having been selected. The input of MaCroDNA has four different settings: `2000_genes_log` is same as the input of Seurat, `all_genes_log` uses the same log-transformation for Seurat but all genes are used, `all_genes_raw` is the original data with all genes, and `noX_genes_raw` is same as the input of clonealign, where the genes on X-chromosome are removed and the values are the original values.

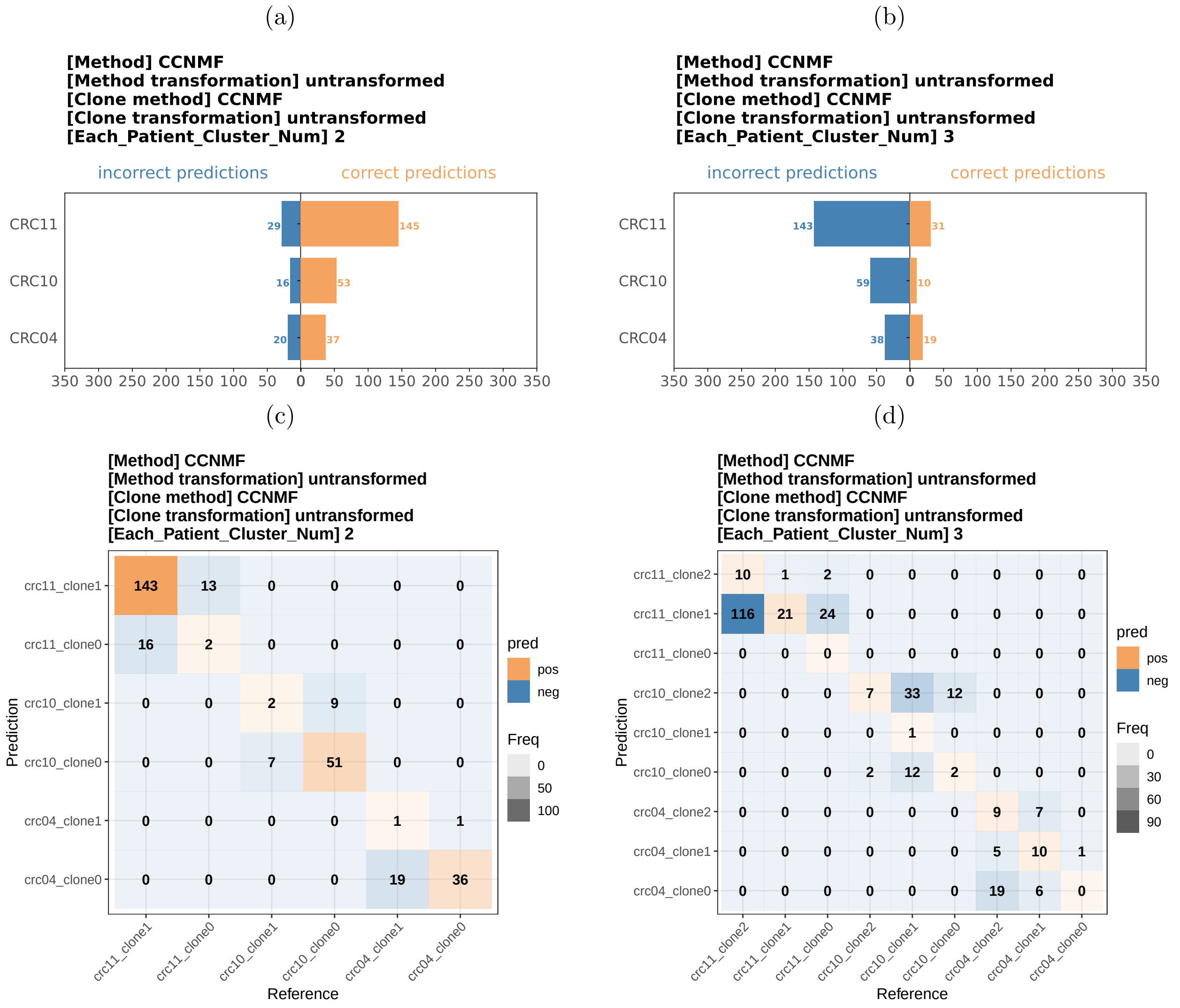


Figure S19: **Results of CCNMF using the original data.** Given a particular number of clusters by the user, CCNMF infers that number of clusters. Here, we performed two experiments, one with two clusters being inferred for each patient, and the other with three clusters for each patient (except for the number of clusters, the rest of CCNMF's input parameters were set as default). The input data are original values. (a) and (c) are results of CCNMF with two clusters in each patient. (b) and (d) are CCNMF's results with three clusters in each patient.

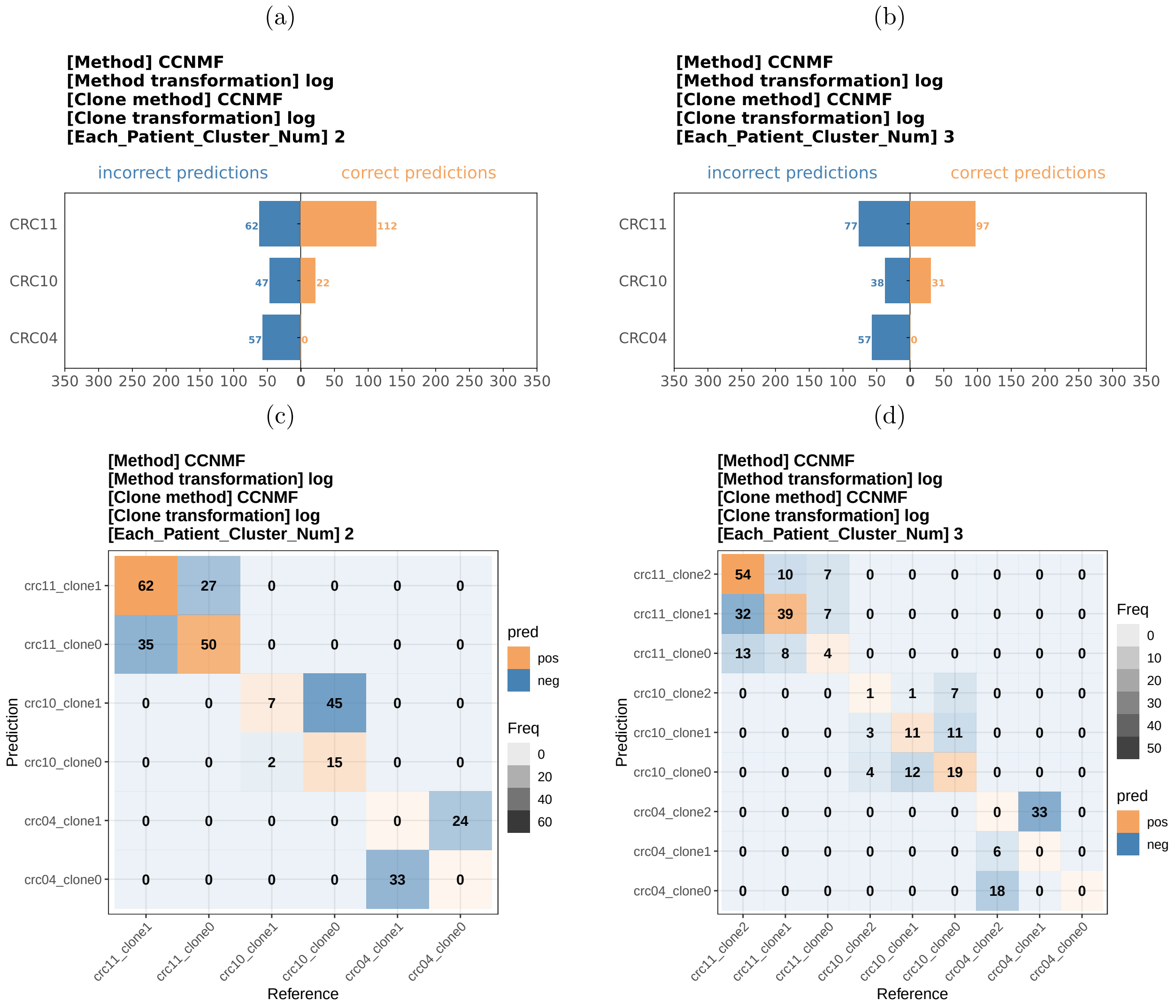


Figure S20: **Results of CCNMF using log-transformed data.** We performed two experiments, one with two clusters being inferred for each patient, and the other with three clusters for each patient. The input data are log-transformed. (a) and (c) are CCNMF's results with two clusters in each patient. (b) and (d) are results of CCNMF with three clusters in each patient.

Table S1: List of the COSMIC genes with statistically significant phylogenetic signal (i.e., $p$-value $<$ 0.05 and $K^* > 1$) identified in the BE data set.

| Patient ID (histology) | COSMIC genes with $p$-value $< 0.05$ and $K^* > 1$[1] |
|---|---|
| 16 (EAC) | *GNAS, BTG1, ATP1A1, B2M, DDX5, H3F3B, MSI2, PABPC1, EIF3E, HSP90AB1, ERBB3, ERBB2, HNRNPA2B1, H3F3A, RPL10, BTG2, MUC1, RPL5, SMARCE1, AKAP9* |
| 20 (HGD1) | *PTPRC, LCP1, IKZF1, CCND2, MSN, PRKCB, TNFAIP3, FLNA, LCK, RHOH, PRF1, BTK, JAK3, PIM1, B2M, FNBP1, NFATC2, ITK, SEPT6, MUC1, ZNF331, PRDM1, SH2B3, IKZF3, GATA2, BLM, ERBB3, BCL2, MYH9, IDH2, LYL1, BCL11B, TAL1, KIAA1549, KIT, MALT1, CYLD, IL7R, CARD11* |
| 14 (HGD) | *CXCR4, MUC16, SLC34A2, AXIN2, ZNRF3, SIRPA, ETV5, HMGA2, TP53, GATA1, SMAD2, CDKN1A, PRF1, CLP1, TFE3, TAL1, BCL2, CBFA2T3, BTK, WAS, ZNF331, PTPRT, FCGR2B, HLF, PTPN13, LYL1, KDSR, FAS, RARA, ITK, VAV1, JAK3, WWTR1, PIM1, SRGAP3, BRAF, PRKCB, POLD1, RMI2, FLI1, IRF4, IKZF1, RBM15, SNX29, CREB3L2, ID3, SEPT6, PRDM1, GATA3, SMAD4, BCL11B, COL1A1, JAZF1, LZTR1, CSF1R, MITF, ZCCHC8, LATS2, PLCG1, DDIT3, RHOH, CD28, CARD11, FNBP1, CDKN2C, NR4A3, ZMYM3, IL7R, HEY1, MNX1, FLNA, BAX, ELF4, MALT1, SMARCB1, SH2B3, PMS2, TBX3, NFATC2, MSN, MAFB, BLM, APOBEC3B, CHST11, STAT5B, ABL2, SMARCD1, CRTC3, SMAD3, XPC, FOXO1, KIT, FOXO4, EXT2, ERCC4, CEP89, CYLD, NCOA2, HOXA11, NOTCH1, TNFRSF14, NTHL1, DGCR8, RABEP1, CCND3, KMT2D, FANCE, SS18, PTPN6, NBEA, ACVR1B, NF2, PTPRC, FIP1L1, TLX1, ATF1, CPEB3, MYCL, CCND2* |
| 6 (HGD) | *ERBB2, SMARCE1, MUC4, MSI2, CD74, KLF6, SUB1, FKBP9, NFIB, PABPC1, MACC1, ETNK1, NACA, TMPRSS2, HIF1A, KIF5B, HMGA1, SF3B1, ALDH2, MUC1, IDH1, ARHGEF10, KIT, HSP90AA1, LCP1, AKAP9, CTNNB1, NPM1, MECOM, MET, EIF4A2, BRD3, CTNNA1, FNBP1, ACSL3* |
| 9 (NDBE) | *ERBB2, LASP1, ARID1B, SF3B1* |
| 20 (CARD) | *MUC1, ALDH2, NDRG1, ELF3, SDC4, KLF4, CCND1* |

[1] The names of the genes are sorted based on their corresponding $K^*$ values in descending order.
